# Propofol overcome Tamoxifen resistance in breast cancer by modulating tumorogenesis, immune response and metabolism

**DOI:** 10.1101/2024.04.15.589641

**Authors:** Runyang Yin, Jing Gao, Chunyan Guo, Yang Liu

**Author notes:** Co-correspondence author. Department of Anesthesiology, The Affiliated Hospital of Inner Mongolia Medical University, Hohhot, Inner Mongolia Province 010050, China. Correspondence author. Address: No. 1 North Passage Road, Huimin District, Hohhot, Inner Mongolia Province, Hohhot, Inner Mongolia Province, China. These authors contributed equally to this work (Runyang Yin, Jing Gao).

## Abstract

Although 70% of estrogen receptor (ER)-positive breast cancer patients benefit from Tamoxifen therapy, the developing resistance to Tamoxifen leads to high rates of metastasis and poor prognosis. Propofol, a commonly used anesthetic, has been shown to inhibit the occurrence and development of breast cancer. However, its role in anti-endocrine resistance in ER-positive breast cancer remains unclear. In this study, we found that Propofol significantly promotes cell cycle arrest, induces apoptosis, and inhibits proliferation in Tamoxifen-resistant breast cancer cells. Furthermore, transcriptome sequencing analysis revealed 1065 differentially expressed genes (DEGs) between Propofol-treated Tamoxifen-resistant MCF-7 (MCF7-TR) cells and none-treated MCF7-TR cells. Gene ontology annotation enrichment analysis (GO), Kyoto Encyclopedia of Genes and Genomes pathway enrichment analysis (KEGG), and gene set enrichment analysis (GSEA) showed that Propofol affects the expression of genes located in the plasma membrane and cell periphery, mainly regulating signals involved in cell cycle regulation, immune response modulation, and metabolism. Our results provide new insights into the mechanism by which Propofol controls resistance to Tamoxifen in breast cancer progression and offer a theoretical basis for clinical treatment.

## 1 Introduction

Breast cancer is one of the most prevalent malignancies in women, accounting for more than 24% of new cancer cases in women worldwide and about 15% of cancer-related deaths, and posing a significant threat to women’s health [18]. In recent years, the incidence and mortality of female breast cancer in the world have shown an increasing trend every year, and the onset tends to be younger individuals[28]. Common treatment approaches for breast cancer include surgery followed chemotherapy or radiotherapy, targeted therapy and immunotherapy, as well as endocrinetherapy. Tamoxifen, as a potent selective estrogen receptor modulator (SERM), has been reported to competitively bind to the estrogen receptor (ER) on tumor cells, thereby preventing the promotion of estrogen on tumor cell growth and proliferation [35]. However, Tamoxifen resistance remains a significant obstacle which lead to tumor relapse and metastasis in most of breast cancer patients, and there is no effective method to reverse Tamoxifen resistance in clinic[6]. Therefore, searching for novel non-toxic, broad-spectrum yet cost-effective alternative agents to overcoming Tamoxifen resistance is crucial against breast cancer.

Propofol is a short-acting anesthetic used to induce sedation during a variety of surgical procedures[15]. A large number of studies have demonstrated that in addition to the sedative hypnotic effect, Propofol also affects the development of tumors, reduces the spread of cancer cells by regulating a variety of signaling pathways, and has direct anti-tumor effects[23; 37; 39]. Propofol was also reported that possessed a protective effect on systemic inflammation and immune function[39]. A multi-center retrospective clinical analysis has suggested that Propofol significantly extends survival after surgery for primary breast cancer compared to sevoflurane anesthesia, which may be related to its ability to reduce the function of breast cancer stem cells, induce apoptosis of breast cancer cells, and inhibit the proliferation of breast cancer cells[8]. Studies have shown that propofol can enhance the sensitivity of breast cancer patients to trastuzumab, reducing the local recurrence of breast cancer patients undergoing conservative breast surgery[30]. All these studies suggest that Propofol may be a promising candidate drug with important bioactivities as a therapeutic strategy against breast cancer. However, the regulation of Propofol on breast cancer resistance to Tamoxifen and its potential mechanisms have not been determined.

The recent progress in transcriptome sequencing has enabled its wider use in identifying targets of antitumor agents against multiple cancers[17; 27]. Based on RNA-seq analysis, novel mechanisms of cancer progression and metastasis involving numerous molecular and signaling pathways, were demonstrated[21]. Accumulating transcriptome sequencing analysis data suggests that tumorogenesis progression (such as cell cycle progression/arrest, apoptosis), immune response, and metabolic regulation play an essential role in the process of drug resistance in breast cancer[25; 32; 40]. It was revealed by transcriptome sequencing analysis that HER2^+^ breast cancer cell lines exposed to Tyrosine Kinase Inhibitor targeting HER2 (HER2-TKIs) produce two Drug tolerant persisters with different transcriptomes, and tumor survival signaling pathways compared to the control [5].

In this study, we found Propofol treatment promoted the cell cycle arrest, induced apoptosis, and inhibited the proliferation in MCF7-TR cells, exerting potential antitumor properties. Further RNA-seq gene expression profiles obtained from Propofol-stimulated MCF7-TR cells were compared with in vitro data, suggesting mutually functional consistency or respectively complimentarity. Taken together, our results showed that Propofol can reverse the resistance to Tamoxifen in breast cancer cells by affecting tumorigenesis, immune response and metabolism regulation. The research will have guiding significance for clinical anesthesia in breast cancer surgery and provide new insights into the prevention and treatment of Tamoxifen-resistant breast cancer.

## 2 Materials and methods

### 2.1 Materials

Propofol and Tamoxifen were purchased from MCE (MedCheExpress, Shanghai, China) company. For the experiment, the Propofol and Tamoxifen were dissolved in DMSO and further diluted in culture medium for storage. (10 mM)

### 2.2 Cell culture and treatments

A human breast adenocarcinoma cell line MCF-7 was bought from American Type Culture Collection (ATCC, Manassas, VA, USA). Cells were grown in complete DMEM media supplemented with 10% FBS and penicillin (100 U/ml) and streptomycin (100 μg/ml) under standard conditions at 37 LJ with 5% CO_2_.

MCF7 cell line resistant to Tamoxifen (MCF7-TR) were modified by treating and keeping the cells alive in media with 5 µg/mL Tamoxifen for 12 weeks to achieve resistance. Comparing the expression of P-gp/ABCB1/MDR1 in Tamoxifen-sensitive parent cell lines to RT-PCR, we could prove that Tamoxifen resistance had developed in the cell lines. Before any experiments were performed, Tamoxifen-free media were set up for 7–10 days.

MCF7-TR cells were seeded onto a 12-well plate with a density of 2 × 10^5^ mL^-1^ and incubated overnight. Propofol was added to the cell culture medium with a final concentration of 5 μM, 10 μM or 20 μM and incubated for 24 h. Control groups were received an equal volume of DMSO. Three repeated wells were included in each group.

### 2.3 Cell cycle and apoptosis detection

The cell cycle and apoptosis were analyzed using a Gallios flow cytometer (Beckman Coulter). Briefly, MCF7-TR cells were seeded at on 12-wells plate with 8×10^4^/mL and incubated overnight. The cells were treated with or without Propofol (20 μM) for 48 hours. Then cell pellets were collected and centrifuged (300g, 5min). For cell cycle analysis, cells were fixed with ice-cold 70% ethanol for for 15min, washed with PBS, and incubated with propidium iodide (PI) staining solution containing 50µg/mL PI, 0.1mg/mL RNaseA, 0.05% Triton X-100 at room temperature for 1 h, and analyzed by Gallios flow cytometry.

Apoptosis detection was performed by FITC Annexin-V/PI commercial kit (Becton Dickenson, Franklin Lakes, NJ, USA) following the manufacture’ instructions. The samples were analyzed by fluorescence-activated cell sorting (FACS) using Gallios flow cytometer (Beckman Coulter) within 1 h after staining. Data were analyzed using Kaluza v. 1.2 (Beckman Coulter).

### 2.4 Cell Viability

Cell viability was assayed using a CCK8 assay kit (Biosharp, China). In brief, MCF7-TR cells were seeded at 6 × 10^3^ cells per well and cultured in 96-well plates overnight. Different concentrations of Propofol were added to each well and then incubated for 48 h. CCK8 solution was then added to each well and incubated for 4 h. The effect of Propofol on cells was measured with absorbance at 490 nm using a spectrophotometer (Synergy HT, Bio-Tek, USA).

### 2.5 Clone formation assay

MCF7-TR cells were seeded in 6-well plates at a density of 1000 cells per well. Then medium was changed every 4 days. On the 6th day, cell clones were photographed using OLYMPUS inverted fluorescence microscope.

### 2.6 RNA extraction, library construction and sequencing

Total RNA was obtained from MCF7-TR cells treated with DMSO or Propofol using TRIZOL reagent according to the manufacture’s protocol (Invitrogen Life Technologies, Shanghai, China). The quality and concentration of the RNAs were detected by Nanodrop spectrophotometer (Thermo Scientific Technology, Shanghai, China). The quality and concentration of the RNA were detected using Nanodrop spectrophotometer (Thermo Scientific Technology, Shanghai, China).

The purified RNA from three repeat samples of solvent control and Propofol-treated MCF7-TR cells was processed to prepare an mRNA-seq library according to Illumina’s standard procedure (protocol no. 15008136)[33]. Briefly, ribosomal RNAs (rRNA) were eliminated by a poly-(A) containing mRNA selection procedure to minimize their sequencing. Then the remaining mRNA were subjected to cDNA strand synthesis, purification, ending-repair, a-tailing-adaptor ligation and polymerase chain reaction (PCR) amplification. Each library was sequenced using an Illumina NextSeq500 instrument (Illumina, San Diego CA, USA).

### 2.7 Functional and signaling pathway enrichment analysis of differentially expressed genes (DEGs)

Data analysis for gene expression were performed based on the Database for functional classification and annotation, visualization, and integrated Discovery. Biological process, cellular component, and molecular function were mapped on each term of the GO database (http://www.geneontology.org/), and signaling pathways were investigated using the Kyoto Encyclopedia of Genes and Genomes (KEGG) (http://www.genome.jp/kegg/).

### 2.8 Gene Set Enrichment Analysis (GSEA)

Gene Set Enrichment Analysis was performed using the GSEA software (https://www.broadinstitute.org/gsea/) with permutation = geneset, metric = Diff_of_classes, metric = weighted, #permutation = 2500.

### 2.9 Statistical Analysis

All the statistical analysis was presented as the mean ± s.e.m (SD) and performed using Prism 7 (GraphPad Software). Statistical significance was evaluated by two-way ANOVA. Results are from three independent experiments performed. Asterisk coding is indicated in Figure legends as: *, *p* < 0.05; **, *p* < 0.01; ***, *p* < 0.001.

## 3 Results

### 3.1 Propofol treatment significantly promotes cell cycle arrest and apoptosis and inhibits proliferation in MCF7-TR cells

To investigate the effect of Propofol on the cell death in MCF7-TR cells, cell cycle progression and cell apoptosis were analysed by flow cytometry. As indicated by Fig. 1A and B, Propofol at the concentration of 10 nM induced a significant increase in cells in the G0/G1 phase and markable decreases in the S as well as the M/G2 phase of cell cycle in MCF7-TR cells compared to control cells, activating the cell cycle arrest. Additionally, Propofol at 20 nM produced induced marked early apoptosis and late apoptosis after 48 h of exposure compared to control untreated cells (**Fig. 1C and D**). Consistent with these findings, Propofol treatment obviously inhibited cell proliferation in a dose-dependent manner, decreasing MCF7-TR cells viability as shown by cell viability assay and Clone formation assay (**Fig. 1E and F**). These results indicated that Propofol induced cell death, exerting potential antitumor properties in resistant breast cancer cells.

**Figure 1.**
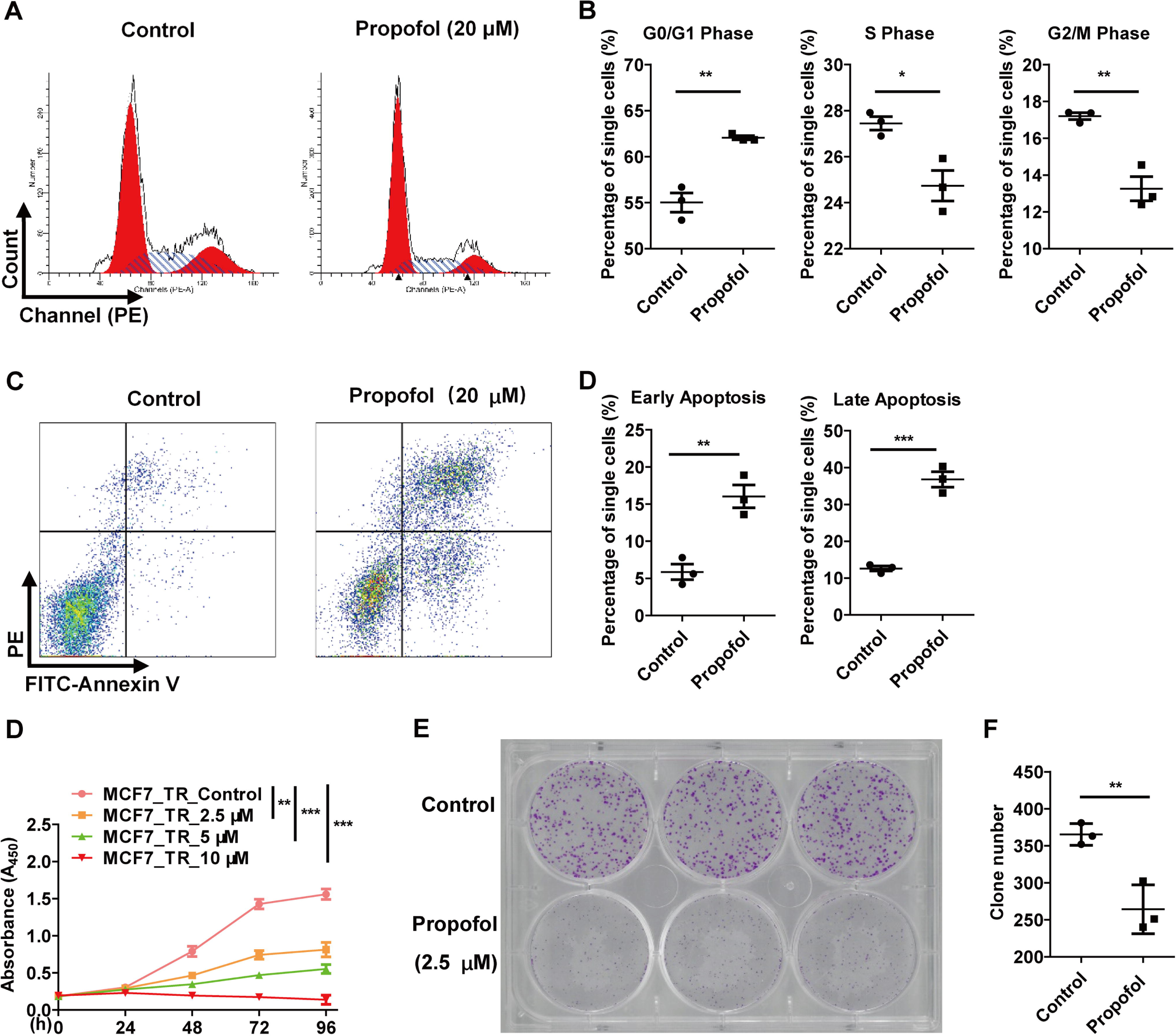
Propofol promotes cell cycle arrest and cell apoptosis in MCF7-TR cells. MCF7-TR cells were treated with or without different concentration of propofol for 24 h. (A) Cell cycle was detected by flow cytometry. (B) Percentage of cells in G0/G1, S and G2/M phase. MCF7-TR cells were treated with or without 20 μM propofol for 24 h. (C) Cell apoptosis was detected by flow cytometry. (D) Percentage of cells in early apoptosis and late apoptosis. Data are shown as the mean ± s.e.m. and are representative of three independent experiments. *, p<0.05, **, p<0.01, ***, p<0.001.

### 3.2 Propofol treatment affects MCF7-TR gene expression profiles

To find out underlying mechanism of Propofol reversed resistance in MCF7-TR cells, we performed transcriptome sequencing analysis for MCF7-TR cells treated with or without Propofol. Principal Component Analysis (PCA) showed good intergroup difference and intragroup consistency between MCF7-TR cells treated with Propofol and without Propofol (**Fig. 2A**).There was a total of 1065 Differentially Expressed Genes (DEGs) in MCF7-TR cells after treatment with Propofol, in which 685 genes were down-regulated and 380 genes were up-regulated dramatically compared with the control (**Fig. 2B, C** and **D**). Overall, Propofol treatment remarkablely altered gene expression profiles in MCF7-TR cells.

**Figure 2.**
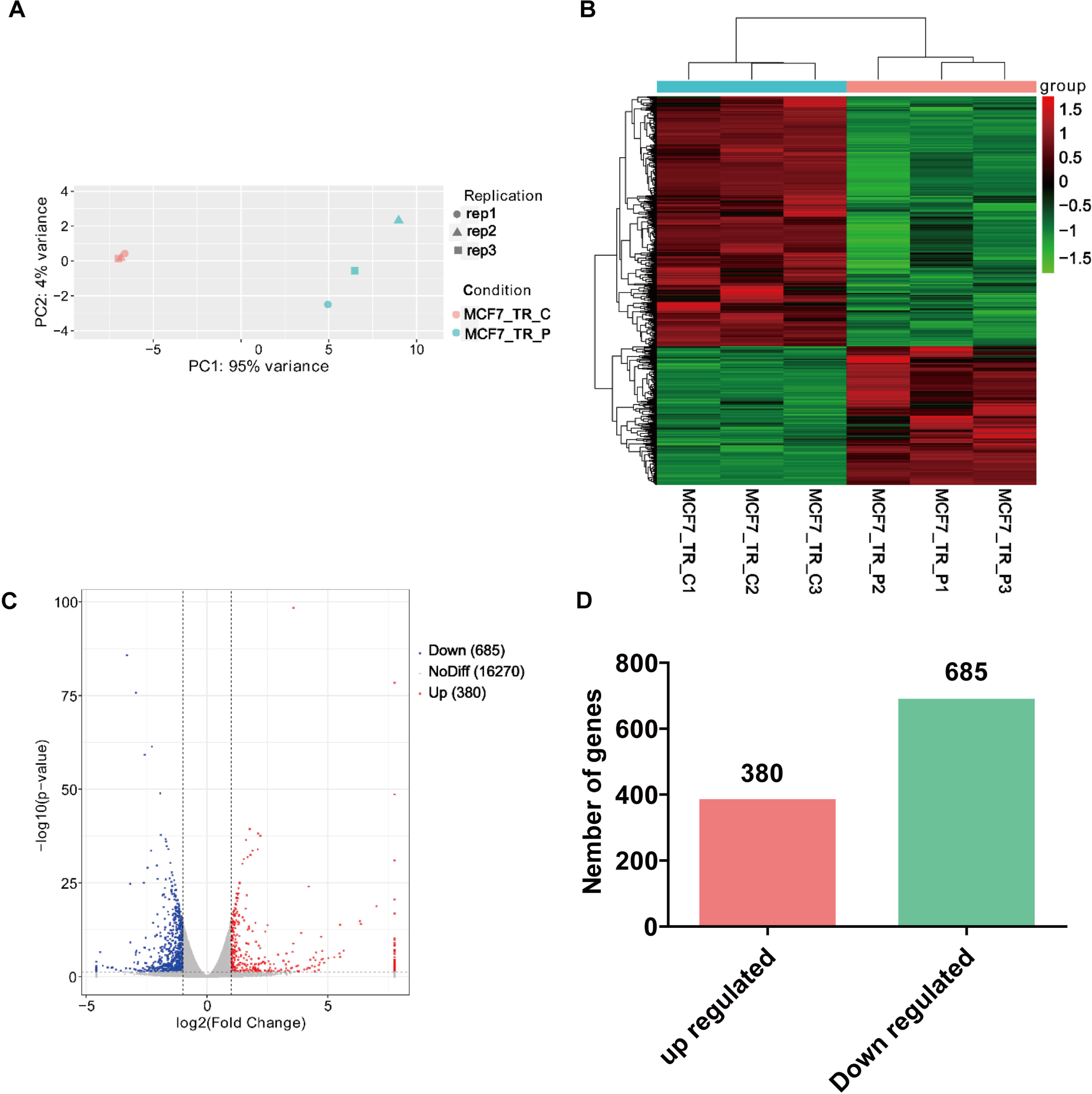
Propofol significantly altered mRNA expression profile in MCF7-TR cells. MCF7-TR cells were treated with or without 10 μM propofol for 24 h. The mRNA expression was detected by RNA-sequencing. (A) PCA analysis shows inter-group differences and intra-group consistency. (B) Hot map shows DEGs between propofol treated or nontreated MCF7-TR cells. (C) Volcano plot shows DEGs between propofol treated or nontreated MCF7-TR cells. (D) Number of DEGs between propofol treated or nontreated MCF7-TR cells. Data are shown as the mean ± s.e.m. MCF7_TR_C1, MCF7_TR_C2 and MCF7_TR_C3 represent the control samples. MCF7_TR_P1, MCF7_TR_P2 and MCF7_TR_P3 represent 10 μM propofol treated samples.

### 3.3 GO terms classification and KEGG pathway enrichment analysis of DEGs in MCF7-TR cells treated by Propofol

In order to unravel the key biological processes that the DEGs were involved in and the molecular function they exerted as well as the pivotal signaling pathways they clustered into in Propofol-treated MCF7-TR cells, GO terms classification and KEGG enrichment analysis were performed. As shown in **Fig.3A**, a large of DEGs were clustered into cell periphery and located in plasma membrane, possessing phosphatase activity and involving in the regulation of immune response and phosphorylation in the MCF7-TR cells upon Propofol exposure. Furthermore, KEGG enrichment analysis showed that many enriched pathways in all treated isolates are related to the process of transcriptional misregulation and cell cycle which occurred in cancer biology. In addition, enriched signaling pathway that were found described more specific were immune reponse processes indicated by cytokine-cytokine receptor interaction and chemokine signaling, and metabolic process indicated by estrogen signaling according to the rich-factor and and FDR values (**Fig.3B**). These results indicated that Propofol treatment could dramatically regulate tumor biology-assosiate process, immune response and metabolism.

**Figure 3.**
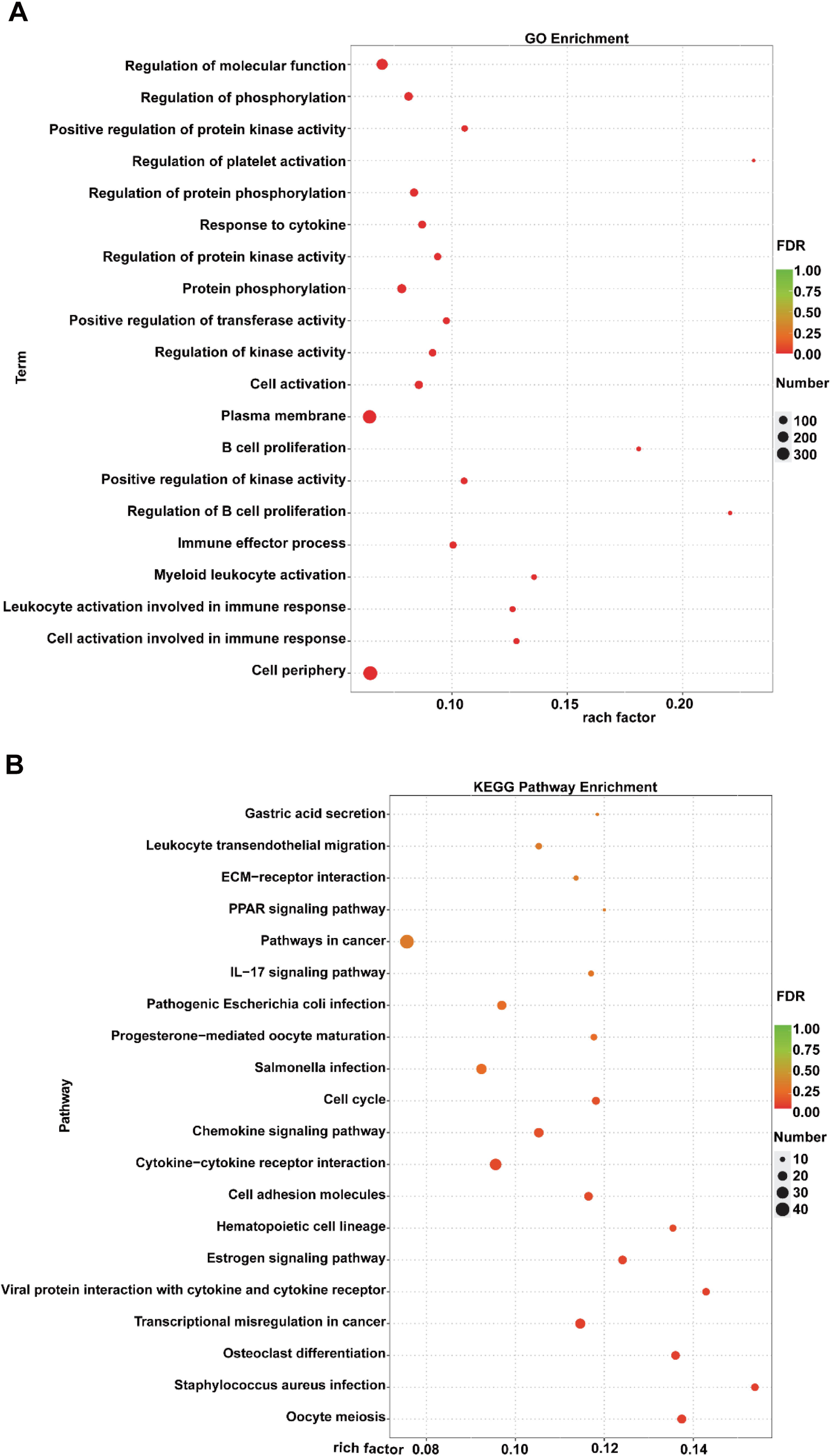
Gene Ontology (GO) terms classification and KEGG pathway enrichment analysis of DEGs in MCF7-TR cells treated by Propofol. Scatter plot for GO annotation analysis of DEGs significantly relevant to biological process, cellular component and molecular function (**A**) and the top 20 most significant KEGG signaling pathways (**B**) obtained from the RNA-sequencing data in MCF7-TR cells after treatment with 10 μM Propofol for 24 h.

### 3.4 Propofol treatment affects gene expression profiles in immune reponse process in MCF7-TR cells

Based on the GO terms classification and KEGG enrichment analysis, we further analyzed the DEGs in immune response-assosiated signaling pathways. As shown in **Fig.4**, the 4 signaling pathways in MCF7-TR cells treated by Propofol were chemokine signaling pathway (**A**), IL-17 signaling pathway (**B**), TNF signaling pathway (**C**), and Leukocyte transendothelial migration (**D**), respectively. Among these pathways, DEGs affected by Propofol treatment were mainly clustered into chemokine signaling pathway, in which 13 genes were upregulated and 7 genes downregulated dramatically. These results suggested that Propofol administration in MCF7-TR cells significantly regulated immune response signals, mainly manifested by chemokine signaling pathway.

**Figure 4.**
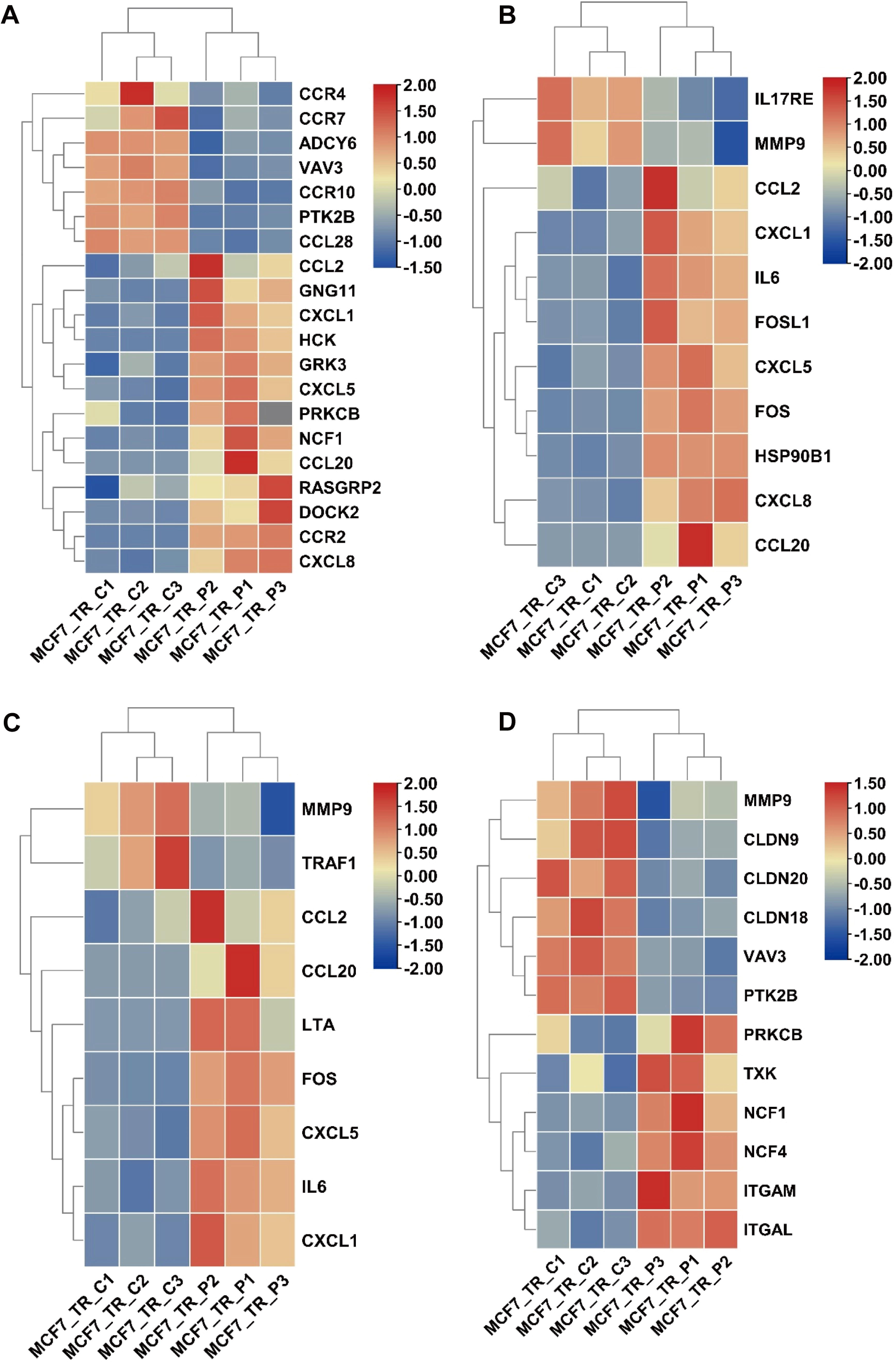
Propofol significantly affected gene expression profiles in immune reponse process in MCF7-TR cells. MCF7-TR cells were treated with or without 10 μM propofol for 24 h. The mRNA expression was detected by RNA-sequencing. (**A**) Hot map shows DEGs in chemokine signaling pathway, (**B**) IL-17 signaling pathway, (**C**) TNF signaling pathway, (**D**) Leukocyte transendothelial migration signaling pathway in MCF7-TR cells treated with or without Propofol.

### 3.5 Propofol treatment affects gene expression profiles in metabolic process in MCF7-TR cells

Next, we analyzed the DEGs enriched obviously in metabolic process indicated in **Fig.5**. The impacts of Propofol on melabolism were characterized by the riboflavin metabolism (**A**), fatty acid biosynthesis (**B**), thyroid hormone synthesis (**C**), and arachidonic acid metabolism (**D**). Among these processes, the effect of Propofol on metabolic regulation is mainly characterized by inhibiting the expression of metabolism-related genes. No up-regulated genes were found in the processes of riboflavin metabolism and fatty acid biosynthesis, and more down-regulated genes were found in the processes of thyroid hormone synthesis and arachidonic acid metabolism in MCF7-TR cells after treatment with Propofol. These findings indicated that Propofol involed in the regulation of metabolic process, mainly depending on the downregulation of the metabolic genes.

**Figure 5.**
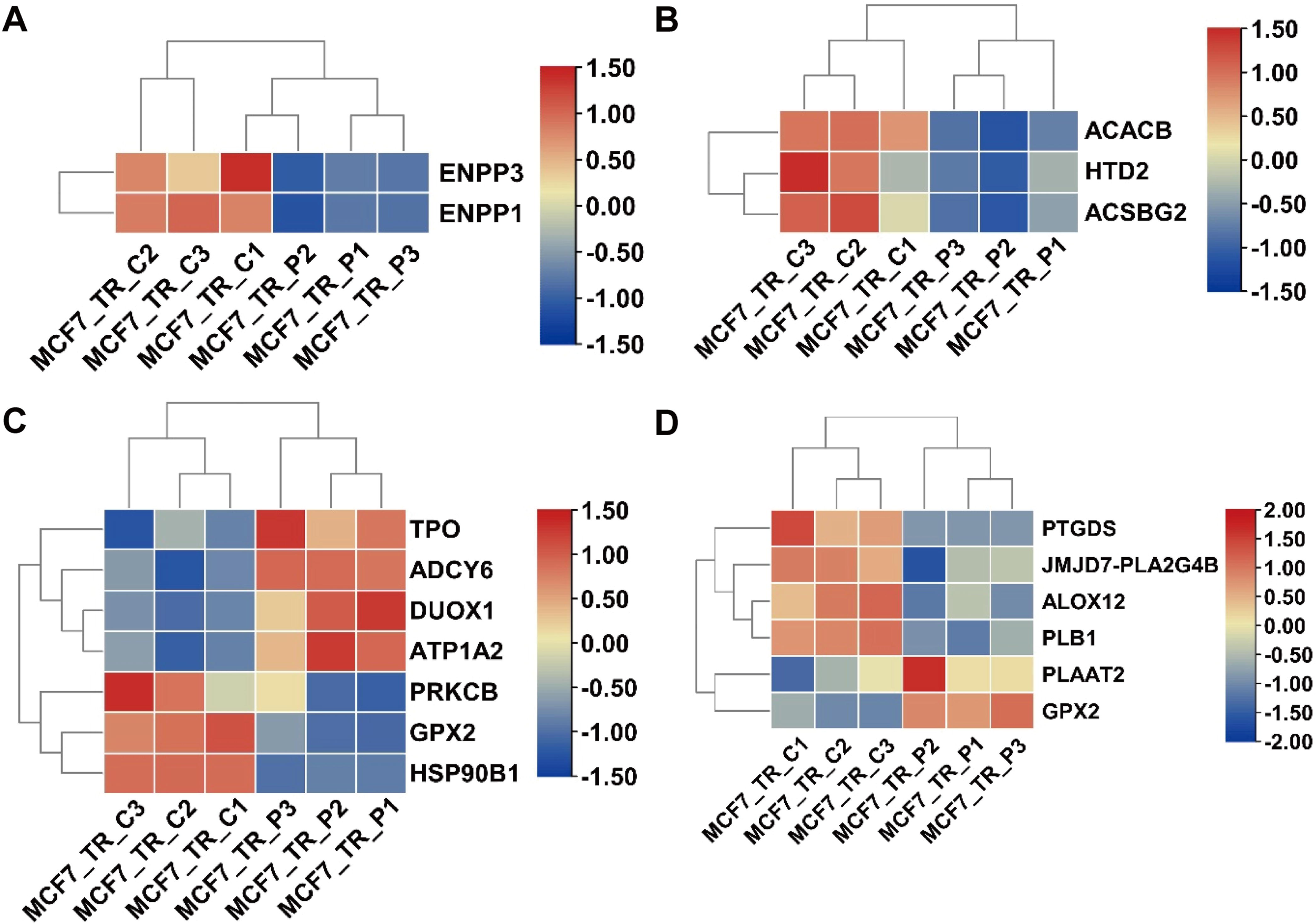
Propofol significantly affected gene expression profiles in metabolism process in MCF7-TR cells. MCF7-TR cells were treated with or without 10 μM propofol for 24 h. The mRNA expression was detected by RNA-sequencing. (**A**) Hot map shows DEGs in riboflavin metabolism process, (**B**) Fatty acid biosynthesis process, (**C**) Thyroid hormone synthesis process, (**D**) Arachidonic acid metabolism process in MCF7-TR cells treated with or without Propofol.

**Figure 6.**
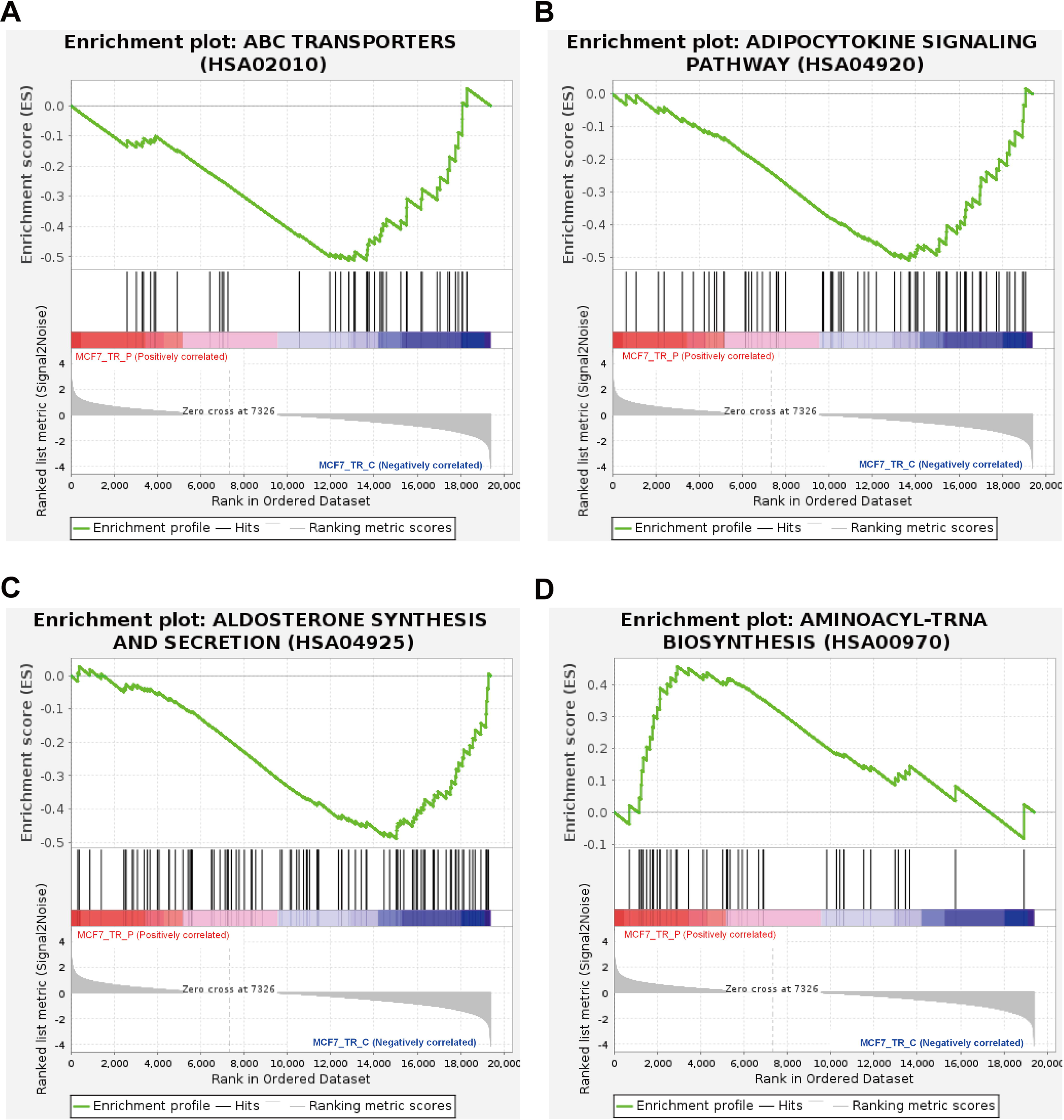
Gene set enrichment analysis (GSEA) of MCF7-TR cells treated with or without Propofol. MCF7-TR cells were treated with or without 10 μM propofol for 24 h. The mRNA expression was detected by RNA-sequencing. (**A**) GSEA plots on Kyoto Encyclopedia of Genes and Genomes (KEGG) pathways in the ABC transporters pathway, (**B**) Adipocytokine signaling pathway, (**C**) Aldosterone sythesis and secretion, (**D**) Aminoacyl-tRNA biosynthesis

### 3.6 Signaling pathways involving in the Propofol sensitivity in MCF7-TR cells

For further characterize the types of molecular functions implicated in immune response and metabolism based on findings in GO and KEGG analysis in Propofol-treated MCF7-TR cells, Gene Set Enrichment Analysis (GSEA) was performed on the RNA-seq data (**Fig.4**). The negatively enriched gene sets involed in ABC transporters (**A**), adipocytokine signaling pathway (**B**), as well as aldosterone sythesis and secretion (**C**), while positively enriched gene set was aminoacyl-tRNA biosynthesis (**D**). These gene sets were composed of genes associated with some metabolites, which indicated that Propofol reversed resistance partially by activating and inhibiting metabolism-related signaling pathways.

## 4 Discussion

Currently, Breast cancer leads in morbidity and mortality among women. Among all breast cancer types, ER^+^ breast cancer is the most common, and ER signaling pathway plays a key role in the occurrence and development of ER^+^ breast cancer[1]. Hence, endocrine therapy that blocks the effects of estrogen has become the first-line treatment for patients with ER^+^ breast cancer[26]. However, the primary and acquired resistance faced by patients receiving endocrine drugs such as Tamoxifen remains an unsolved clinical challenge. Our study demonstrated that Propofol can affect cell cycle progression and induce cell apoptosis in breast cancer Tamoxifen-resistant cell lines through altering the gene expression profiles involving in the regulation to the tumorigenesis, immune response and metabolism.

Much of the research suggested that the development, metastasis and recurrence of tumors are closely related to the cell cycle, apoptosis and proliferation, and the disorders in these events have been confirmed to be the fundamental hallmarks in human malignancies[2; 13; 16]. Indeed, the misregulation in cell cycle related siganling pathways results in the unlimited proliferation of tumour cells. Moreover, it was reported that one of the approaches for increasing the sensitivity of cancer cells to chemotherapeutics was to in synergy them with cell cycle regulators [24].Preclinical studies have shown that Propofol can significantly inhibit the development of breast cancer tumors[29; 31; 36]. In this study, the detected anti-tumoural activity of Propofol was characterised by inhibited cell cycle progression, increased cell aproposis, in parallel with the inhibited proliferation, indicating that Propofol seems to act as a cell cycle regulator to overcome resistance to Tamoxifen in breast cancer cells.

In concordance with the *in vitro* results, data obtained from transcriptome sequencing analysis showed that a large of DEGs in MCF7-TR cells treated with Propofol were enriched into the process of cell cycle and transcriptional misregulation, which were in correlation with Propofol-counteracted resistance to Tamoxifen. Cell cycle mainly consist of two key events – interphase and the mitotic or M phase. Transitions between the cell cycle phases are triggered by the cyclin-dependent kinases (CDKs) and their binding lingands in the cyclin proteins family[10; 20]. Cyclin-dependent kinase 4/6 (CDK4/6) inhibitors have been employed as potent, selective and orally bioavailable treatments for hormone receptor-positive (HR+), human epidermal growth factor receptor 2-negative (HER2^−^) breast cancer[7; 9]. Currently, each of CDK4/6 inhibitors including Ribociclib, Palbociclib, and Abemaciclib were reported to combined with endocrine therapy as the first-line or second-line therapies, showing the significant improvements in progression-free survival (PFS)[14]. It have been elucidated that CDK4/6 inhibitor can induce the G1/S phase cell cycle arrest through selectively targeting CDK4/6, which control cell cycle progression via retinoblastoma protein phosphorylation[3]. Propofol administration in our results showed a remarkable inhibition to S phase of cell cycle in MCF7-TR cells, implying a potential correlation with CDK4/6 signals. Preclinical studies showed many cell cycle-assosiated proteins were implicated in CDK4/6 inhibitors’ resistance. CDK4/6 inhibition and cytostasis evasion are two critical events occurred in ER-positive breast cancer cells, and combined targeting of both CDK4/6 and PI3K can induce cancer cell apoptosis in vitro and in patient-derived tumor xenograft (PDX) models, and prevent resistance to CDK4/6 inhibitors by reducing the levels of cyclin D1 (CCND1) and other G1-S cyclins, ultimately resulting in tumor regression[12]. The above studies indicate that the targets of Propofol reversing resistance to Tamoxifen in breast cancer cell may also act as a part of G1-S cyclins, which need to be explored further in the future.

In addition to the DEGs enriched in the cell cycle involving in the regulation of tumor occurrence and recurrence, those affected by Propofol treatment were also clustered into immune and metabolic signaling pathways. Our results from KEGG pathway analysis indicated that there are some DEGs significantly enriched into chemokine signaling pathway. Research showed that C-X-C motif ligand 10 (CXCL10), as a pro-inflammatory cytokine secreted by tumor cells, played vital role in Tamoxifen resistance in breast cancer[19]. CXCL10 promoted breast cancer tumor proliferation and growth via both estrogen-dependent and independent pathways, while CXCL10 inhibition reversed the resistance of cells to Tamoxifen[34]. Beyond that chemokines and its receptors such as CX3CL1, CXCL2 and CXC-4 were also identified as the biomarkers for Tamoxifen-resistant therapy[4; 11]. Moreover, our results showed almost 40 DEGs contribute to the regulation in cytokine-cytokine receptor interaction, manifested by the activation of IL-17 signaling pathway and TNF signaling pathway. TRAF4 is the member of tumor necrosis factor receptor-associated factor (TRAF) family, which functions as a signal transducer in TNF signaling pathway[22]. It was shown that TRAF4 was overexpressed in Tamoxifen-resistant breast cancer cell line, and knockdown of TRAF4 can overcome the resistance to Tamoxifen[38]. In terms of our results, TRAF4 may function as potential target in Propofol-treated MCF7-TR cells, which need to be elucidated further. In addition, metabolic processes such as Riboflavin metabolism, Fatty acid biosynthesis, Thyroid hormone synthesis and Arachidonic acid metabolism in MCF7-TR cells were response to Propofol in the current study, in which most genes’ expression were downregulated. Kim et al. found that deregulation of a gene in fatty acid metabolism can cause drug resistance[19]. Furthermore, results from GSEA analysis also showed that significant metabolites including adipocytokine, aldosterone and aminoacyl-tRNA were strongly correlated with Propofol sensitivity, suggesting the activation and/or inhibition of the signaling pathways they involved in played an important role in Propofol-counteracted resistance to Tamoxifen. Collectivelly, the antitumor activities of Propofol have a strong connection with its impacts on immune response and metabolism according to the study.

In conclusion, Propofol contributes to promoting cell cycle arrest, triggering apoptosis and repressing proliferation, reversing resistance to Tamoxifen in breast cancer cells, possessing the potential to be utilized as a cell cycle regulator for the treatment of breast cancer. Specially, deep investigations on the DEGs affected by Propofol in cell cycle process, immune and metabolic signaling pathways are urgently needed.

## Data availability

The datasets produced in this study are available in the following databases: NCBI BioProject with the accession number: PRJNA1021154. (https://www.ncbi.nlm.nih.gov/bioproject/PRJNA1021154).

## Acknowledgement

This work was supported by grant from Scientific research project of universities in Inner Mongolia Autonomous Region (NJZZ23016).

## CrediT Contribution Statement

Conceptualization: Yang Liu, Runyang Yin and Jing Gao; Formal Analysis: Runyang Yin, Jing Gao and Chunyan Guo; Writing original draft: Runyang Yin and Jing Gao.

## Declaration of competing interest

There are no conflicts to declare.

## Notes

https://www.ncbi.nlm.nih.gov/bioproject/PRJNA1021154

### Competing Interest Statement

The authors have declared no competing interest.

